# Myelinated fiber labeling and orientation mapping of the human brain with light-sheet fluorescence microscopy

**DOI:** 10.1101/2025.03.31.645981

**Authors:** Michele Sorelli, Danila Di Meo, Samuel Bradley, Franco Cheli, Josephine Ramazzotti, Laura Perego, Christophe Destrieux, Patrick R. Hof, Francesco S. Pavone, Giacomo Mazzamuto, Irene Costantini

## Abstract

The convoluted network of myelinated fibers that supports behavior, cognition, and sensory processing in the human brain is the source of its extraordinary complexity. Advancements in tissue optical clearing, 3D fluorescence microscopy, and automated image analysis have enabled unprecedented insights into the architecture of these networks. Here, we investigate the multiscale organization of myelinated fibers in human brain tissue from the brainstem, Broca’s area, hippocampus, and primary visual cortex by exploiting a specific fiber staining method, light-sheet fluorescence microscopy (LSFM), and an advanced spatial orientation analysis tool. Using an optimized protocol that integrates tissue clearing with the lipophilic DiD probe to achieve uniform and deep myelinated fiber labeling, we generate micrometer-resolution volumetric reconstructions of multiple brain regions through an inverted LSFM. Automated image processing, employing unsupervised 3D multiscale Frangi filters, provides orientation distribution functions and local orientation dispersion maps. This enables precise characterization of the directionality of white matter bundles, linking mesoscopic structural properties to orientation details computed at the native micrometric resolution of the LSFM apparatus. The presented workflow illustrates a robust platform for large-scale, high-resolution brain mapping, which may facilitate the investigation of pathological alterations with unparalleled spatial resolution and, furthermore, the validation of other neuroimaging modalities.

## Introduction

The complexity of the brain arises from its intricate network of connections, which underpin cognition, behavior, and sensory processing. Among the key components of brain structure, myelinated fibers play a pivotal role in ensuring efficient neural communication across the central nervous system. Three-dimensional (3D) mapping of this intricate network can provide insights into how the brain integrates information across different regions to produce coherent outputs. It can also facilitate the correlation of specific brain regions with their functional roles, enhancing our understanding of how various areas contribute to cognitive processes and behaviors, and how their structural alteration may have an impact on neurological disorders. Over the years, a variety of methods have been developed to stain and visualize myelinated fibers, each offering unique advantages and limitations. Traditional histological techniques, such as Sudan Black or Luxol Fast Blue, provide high-resolution insights into the microstructural organization but are labor-intensive and restricted to two-dimensional (2D) applications [1, 2, 3]. Immunohistochemical techniques, using antibodies that specifically bind key structural components of myelin, offer high specificity but their large molecular size hinders their diffusion, resulting in penetration difficulties in thick tissues [4, 5, 6, 7]. Label-free approaches that leverage endogenous contrast or autofluorescence, such as coherent anti-Stokes Raman scattering (CARS) [8, 9], third-harmonic generation microscopy (THG) [10], or two-photon fluorescence microscopy (TPFM) with MAGIC (Myelin Autofluorescence imaging by Glycerol Induced Contrast enhancement) [11, 12], can also be used. Nonetheless, these techniques possess limitations regarding imaging penetration depth, acquisition speed and the total tissue volume that can be analyzed. Recent advancements in tissue optical clearing and volumetric imaging techniques, such as light-sheet fluorescence microscopy (LSFM), have enabled unprecedented imaging depth (mm range) and resolution (➭m range) in brain tissue studies, leading to detailed visualization and distinction of complex structures, such as the architecture of myelinated fibers [13, 14, 15, 16, 17, 18]. This can be especially valuable for providing the required ground truth orientation information for the fine quantitative validation of diffusion-weighted magnetic resonance imaging (dMRI) which, despite being already widely used in clinical settings for in vivo white matter imaging, is nowadays still suffering from a maximum achievable spatial resolution of ≈1 mm [19]. Other advanced optical modalities relying on the myelin sheath birefringence, such as polarized light imaging (PLI) [20], are instead restricted to 50-100 ➭m-thick tissue slices, which limits the possibility of accurately quantifying the through-plane inclination of nerve fiber bundles. Automatic serial sectioning polarization-sensitive optical coherence tomography (PS-OCT) [21, 22], on the other hand, shows promising potential for mapping microscopic fiber orientation in large-scale samples, such as whole coronal sections of a human brain hemisphere. However, its spatial resolution cannot currently match the one achieved by 3D fluorescence microscopy, being still limited to ≈10 ➭m [23]. In addition, the applicability of those techniques is limited to connectomic evaluation, whereas clearing methods combined with LSFM imaging enable the characterization of human tissues at the cellular level through specific staining mediated by antibody-epitope recognition [24, 25, 26, 27, 28].

Within the realm of 3D fluorescence microscopy, small fluorescently labeled lipophilic probes can be powerful tools for generating sharp volumetric reconstructions of the myeloarchitecture [1, 24, 29, 30, 31, 32, 33]. While they exhibit an affinity for lipid-rich structures and stain myelin sheats through accumulation rather than specific protein binding, their small size offers an advantage for rapid and deep penetration in large brain tissue volumes. Indeed, when combined with tissue-clearing methods and high-throughput LSFM, they can provide powerful capabilities for generating sizable, high-resolution image datasets, creating a robust platform for large-scale detailed brain mapping studies. The combination of fiber staining and cell-specific identification in cleared human tissues can therefore give a comprehensive understanding of the brain organization, by integrating fiber connectivity information with cytoarchitectural insights at the microscale level. In this regard, automated image processing techniques are essential for the analysis of large 3D microscopy datasets, as they enable efficient, consistent, and objective evaluation of complex biological structures, allowing 3D microscopy modalities to evolve from simple observational to valuable quantitative tools [34, 35, 36, 37]. Manually processing the vast amounts of data generated by LSFM would be not only extremely time-consuming but also prone to inconsistencies and human error. By automating LSFM image processing, researchers can reduce the image analysis time bottleneck, accelerating the overall workflow while maintaining high accuracy and reproducibility. This capability is crucial for advancing scientific understanding, especially in neurobiology, where large high-quality datasets are key to uncovering new insights.

In this work, we develop a specific fiber staining method compatible with tissue clearing and a custom-engineered analysis tool to investigate the multiscale 3D organization of myelinated fibers of different human brain tissues — namely, the brainstem, Broca’s area, hippocampus, and primary visual cortex. The homogenous and deep staining of myelinated fibers using the lipophilic DiD probe was obtained in combination with the SHORT clearing method [24, 25] and immunofluorescence staining of different markers. Micrometer-resolution mesoscopic image reconstructions (in the centimeter range) of the four anatomical brain regions obtained with a custom-built LSFM setup [27, 38] were analyzed using Foa3D, an unsupervised image processing tool built around 3D multiscale Frangi filters, previously used on TPFM images [11]. Here, this tool was optimized to facilitate the analysis of larger tissue reconstructions, such as those obtained with LSFM imaging. This was achieved by reducing memory footprint and leveraging GPU-accelerated image filtering for ensuring that overall image processing times remained substantially shorter than image acquisition times. This demonstrated the capability to reliably detect and characterize the orientation of 3D fiber structures in grey and white matter at the native resolution of the LSFM apparatus employed, without placing a significant bottleneck after imaging.

Overall, the workflow described in the present work — including myelinated fibers labeling in optically cleared tissue, LSFM imaging, and fast automated 3D analysis of fiber orientations — enables an unprecedentedly detailed understanding of the myeloarchitecture of human brain tissue in physiological and pathological conditions. This approach paves the way for the validation of other neuroimaging modalities, such as the assessment of the meso- and macro-scale brain connectivity information targeted by state-of-the-art dMRI, and facilitates the investigation of abnormal states with unsurpassed spatial resolution in large-scale brain microscopy datasets.

## Results

### DiD staining of SHORT-cleared human brain samples

Lipophilic probes that intercalate into membrane phospholipids are commonly used to trace myelinated fibers, but they may not always be compatible with clearing protocols. We used the SHORT method for tissue clearing and antibody labeling and optimized a dedicated fiber staining method employing the lipophilic DiD probe. The synergy of these approaches enabled the 3D reconstruction of both the architecture of neuronal bodies and their connections. Human brain samples were first processed using the SHORT protocol, as previously described [24, 25], for lipids removal and antibody-mediated labeling of different classes of neurons. Myelinated fiber staining was then performed as follows, prior to refractive index (RI) matching in 2,2✬-Thiodiethanol (TDE). SHORT-processed slabs were first equilibrated in a solution containing 10 mM sodium dodecyl sulfate (SDS) for 24 h. DiD was then applied to the slabs through two consecutive incubations in different solutions, to enhance tissue permeability: 0.025 mg/mL DiD in 10 mM SDS (D-off solution) and 0.025 mg/mL DiD in 1X PBS with 0.1% TritonX-100 (D-on). The D-off solution, containing SDS, slowed the incorporation of the dye into membranes, allowing DiD molecules to disperse uniformly throughout the tissue depth. The D-on solution facilitated the integration of DiD molecules into the myelin sheath, enabling a selective visualization of myelinated fibers (Fig. 1). Under the confocal spinning disk microscope at higher magnification, we observed that our DiD protocol enabled the distinct visualization of the myelin-rich edges of single axons, oriented differently throughout the tissue slab (Fig. 1b). At lower magnification, we obtained a detailed imaging of myelinated fiber bundles in different areas of the brainstem, including the pons (Fig. 1c) and midbrain (Fig. 1d). Our experiments revealed that successfully labeling both myelinated fibers and neuronal bodies requires incubating tissue slabs with antibody solutions first, followed by the application of the DiD probe. When the order was reversed — applying the DiD probe before antibody incubation — the pan-neuronal marker NeuN failed to produce a specific signal. Moreover, we observed that increasing the incubation time in either solution (D-off and D-on) did not improve DiD molecule penetration; in fact, it made it worse. Similarly, increasing DiD dye concentrations or reducing the incubation temperature to 4 ^◦^C instead of 37 ^◦^C, did not enhance the dye penetration inside the tissue slabs. With this optimized protocol, DiD can be used as an effective probe for labeling myelinated fibers and can be successfully employed for deep-brain imaging in conjunction with immunostaining of neuronal bodies, enabling high-resolution 3D reconstruction of the human neural architecture.

**Figure 1:**
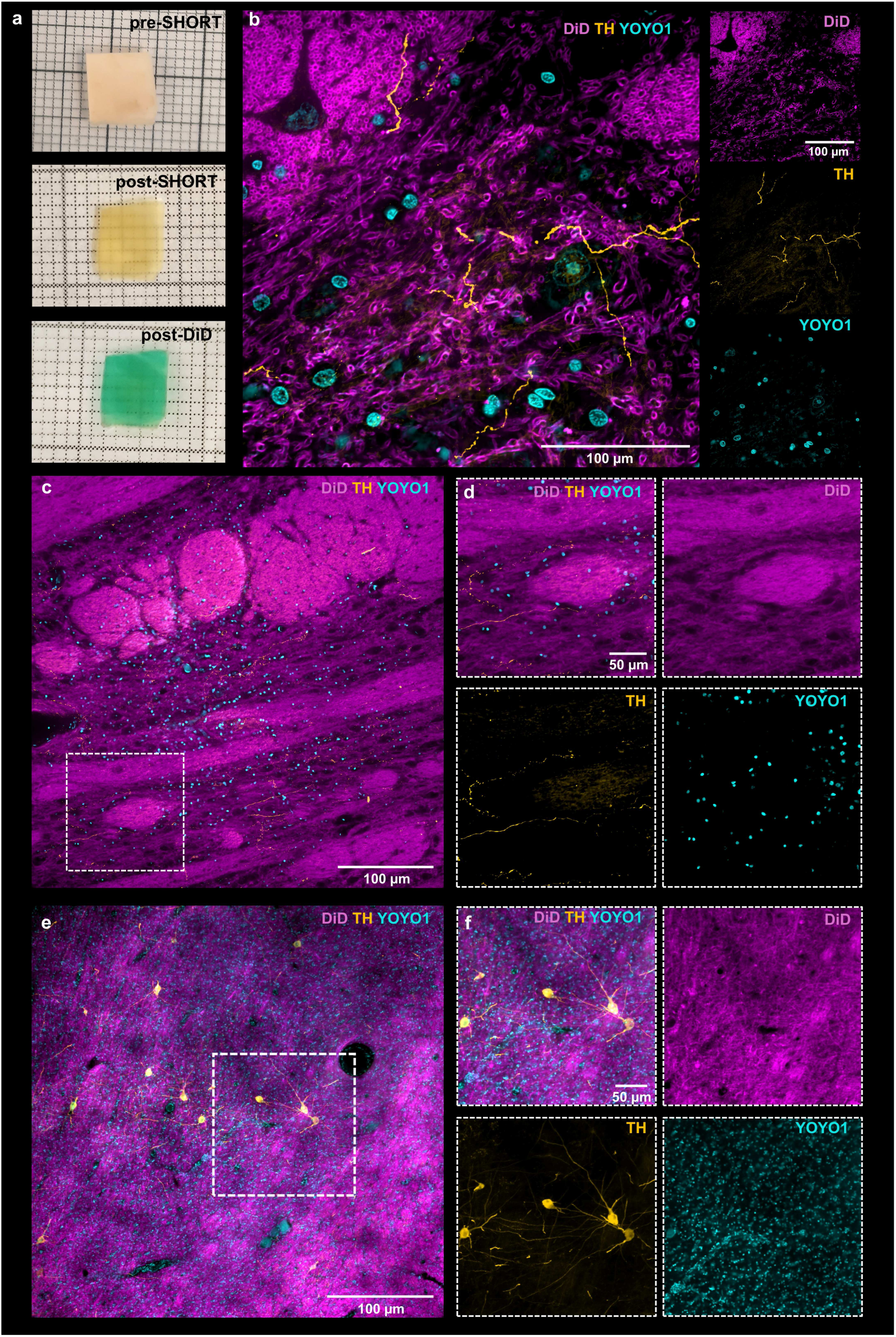
Co-labeling of neurons and myelinated fibers stained with DiD in the human brain. a) Images showing a portion of a brainstem slab (thickness: 300 ➭m) before (pre-SHORT) and after (post-SHORT) clearing and labeling with the SHORT method and after myelinated fiber labeling with the DiD probe (post-DiD). b) Confocal spinning disk images of a SHORT-processed human brainstem slab (pons) labeled with DiD (magenta), an antibody against TH (yellow), and YOYO1 (cyan). Objective: 60X/1.4NA (voxel size: 0.11 ➭m X 0.11 ➭m X 1 ➭m). Single channel images are shown on the right. c) Confocal spinning disk images of a SHORT-processed human brainstem slab (pons) labeled as in (b). Objective: 20X/0.75NA (voxel size: 0.32 ➭m X 0.32 ➭m X 2 ➭m). d) Magnified inset images of (c). e) Confocal spinning disk images of a SHORT-processed human brainstem slab (midbrain) labeled as in Objective: 10X/0.5NA (voxel size: 0.65 ➭m X 0.65 ➭m X 2 ➭m). f) Magnified inset images of (e).

### Volumetric reconstruction of myelinated fibers and cytoarchitectural structure in human brain areas

Adult postmortem human brain tissue slabs, ranging from 300 ➭m to 500 ➭m in thickness, were treated as previously outlined, undergoing SHORT-mediated clearing and antibody labeling, along with selective fiber staining using the DiD probe (Fig. 2). Volumetric imaging was performed using a custom-built dual-view inverted light-sheet fluorescence microscope, achieving a final voxel size of 3.6 ➭m X 3.6 ➭m X 3.6 ➭m, following postprocessing [26, 38].

**Figure 2:**
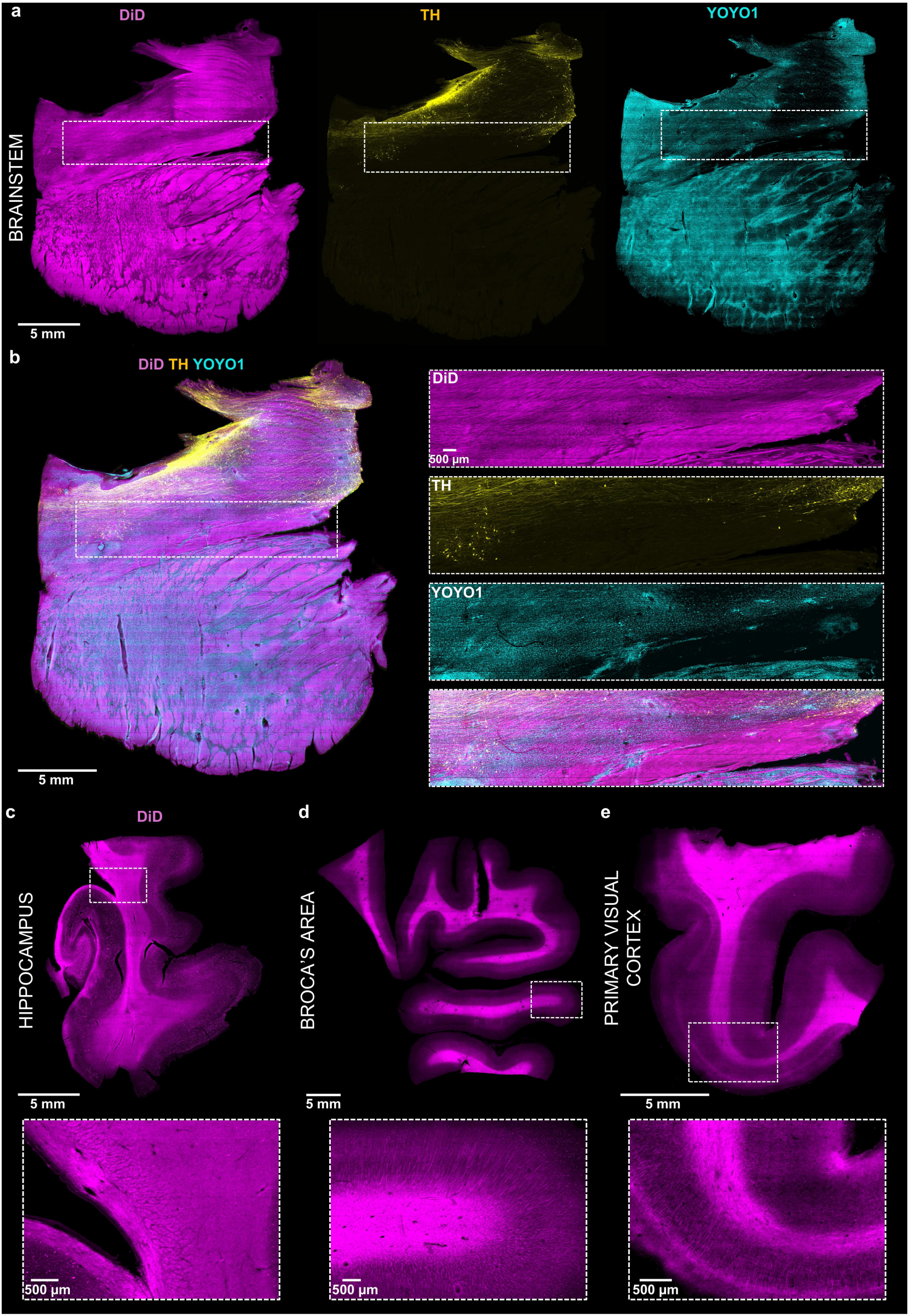
Co-labeling and LSFM imaging of myelinated fibers in different human brain areas. a) Maximum intensity projection (MIP) images (voxel size: 3.6 ➭m X 3.6 ➭m X 3.6 ➭m) of a SHORT-processed human brainstem slab (thickness: 300 ➭m) labeled for DiD (magenta), TH (yellow) and YOYO1 (cyan). b) The multichannel MIP is shown from (a). The magnified inset images on the right highlight regions of interest in the white boxes. c-d-e) MIP images of 500 ➭m-thick SHORTprocessed slabs: (c) postmortem human hippocampus (d) Broca’s area and (d) primary visual cortex, all labeled with the DiD probe. White boxes correspond to the magnified inset images at the bottom.

The slab from a brainstem region (pons) showed uniform multi-labeling with an antibody against tyrosine hydroxylase (TH), enabling sparse labeling of individual noradrenergic neurons and axons, mainly localized in the locus coeruleus. This was combined with YOYO1 staining for cytoarchitectural definition of different neuronal nuclei (Fig. 2a, 2b and S1a). DiD-labeled myelinated fiber bundles of different thicknesses and organization were clearly visible throughout the depth of the 3D tissue reconstruction (Fig. 2a, 2b, and S1a).

Volumetric reconstruction of slabs from human Broca’s area (Fig. 2c, S1a), hippocampus (Fig. 2d, S1a) and primary visual cortex (Fig. 2d, S1a) revealed densely packed myelinated fiber bundles (i.e., white matter), with radial myelinated axons bundles branching from the innermost layer of the grey matter, where neuronal bodies were present. Moreover, neuronal immunostaining was also performed with antibodies against somatostatin (SST) and calretinin (CR), to visualize distinct neuronal subtypes and against NeuN, for the detection of all neurons (Fig. S2a, S2b and S2c).

A slab from the postmortem human brainstem was also cleared using the organic solvent-based method iDISCO [39] and subjected to the DiD staining protocol, as previously described. Although the dye appeared to penetrate deeply into the tissue that assumed the characteristic blue appearance, after delipidation with 100% dichloromethane (DCM) the blue color was lost and LSFM imaging showed the absence of a specific fluorescence signal (Fig. S1b). Thus, a subset of lipids is likely preserved in human tissues processed with the SHORT method, enabling deep visualization of the multilayered, lipid-rich myelin sheath together with multiple subpopulations of neurons in different human brain tissues, an outcome not achieved with the iDISCO method.

### Foa3D for LSFM image analysis

The original version of Foa3D (*Fiber orientation analysis (Foa3D)* in *Materials and Methods*) was designed to iteratively slice and process basic sub-volumes of tiled TPFM tissue reconstructions [11]. Although optimized to utilize the full available memory, this approach was limited to a single CPU core. This limitation could become particularly significant in terms of processing time when analyzing the larger volumetric image reconstructions which are typically acquired using LSFM setups (≈10-1000 GB). Therefore, to enhance efficiency, whenever possible, the improved Foa3D tool leverages a GPU-accelerated Frangi filtering stage — the unsupervised fiber enhancement stage — that typically constitutes the primary bottleneck in the fiber orientation analysis pipeline. This optimization significantly reduces overall image processing times, thereby streamlining brain tissue myeloarchitectural studies. The computational cost of the two implementations reported above is summarized in Table 1. The new GPU-accelerated configuration demonstrates a remarkable and consistent decrease in execution time, achieving reductions of approximately 98% across all regions. This suggests that the implementation effectively leverages the GPU’s massive parallel processing capabilities, effectively neutralizing the image processing bottleneck following LSFM tissue imaging.

**Table 1:** Computational cost of two configurations of Foa3D’s Frangi-filter-based multiscale unsupervised enhancement of myelinated fibers (Frangi filter scales: [5, 10] ➭m; ODF scales: [250, 1000] ➭m). The new GPU-accelerated configuration demonstrates a remarkable and consistent decrease in processing time, achieving reductions of ≈98% across all regions.

| Human<br>Brain<br>ROI | Image Size |  | Time [h:min:s] |  | ODF<br>CPU |
| --- | --- | --- | --- | --- | --- |
|  | [GB]<sup>†</sup> | [mm<sup>3</sup>] | Frangi filter<br>CPU | GPU |  |
| Brainstem | 25.4 | 636.7 | 21:45:29 | 0:25:04 | 0:37:42 |
| Broca’s area | 38.1 | 954.8 | 24:45:36 | 0:33:06 | 0:52:12 |
| Primary visual cortex | 7.7 | 194.1 | 6:53:19 | 0:09:09 | 0:10:48 |
| Hippocampus | 14.5 | 363.8 | 13:16:08 | 0:19:28 | 0:10:30 |

<sup>†</sup> *uncompressed*

### 3D myelinated fiber orientation

Fig. 3 shows myelinated fiber orientation images generated using Foa3D from the four distinct anatomical regions of the human brain — brainstem, Broca’s area, hippocampus, and primary visual cortex — imaged with LSFM. The high-resolution spatial orientation Hue-Saturation-Value (HSV) colormaps (Fig. 3a-d, voxel size: 3.6 ➭m X 3.6 ➭m X 3.6 ➭m) emphasize the capability of the Foa3D tool to correctly describe the myeloarchitecture of the brain tissue samples, resolving the organization of intricate fiber structures without sacrificing the micrometer resolution achieved by the employed LSFM apparatus.

**Figure 3:**
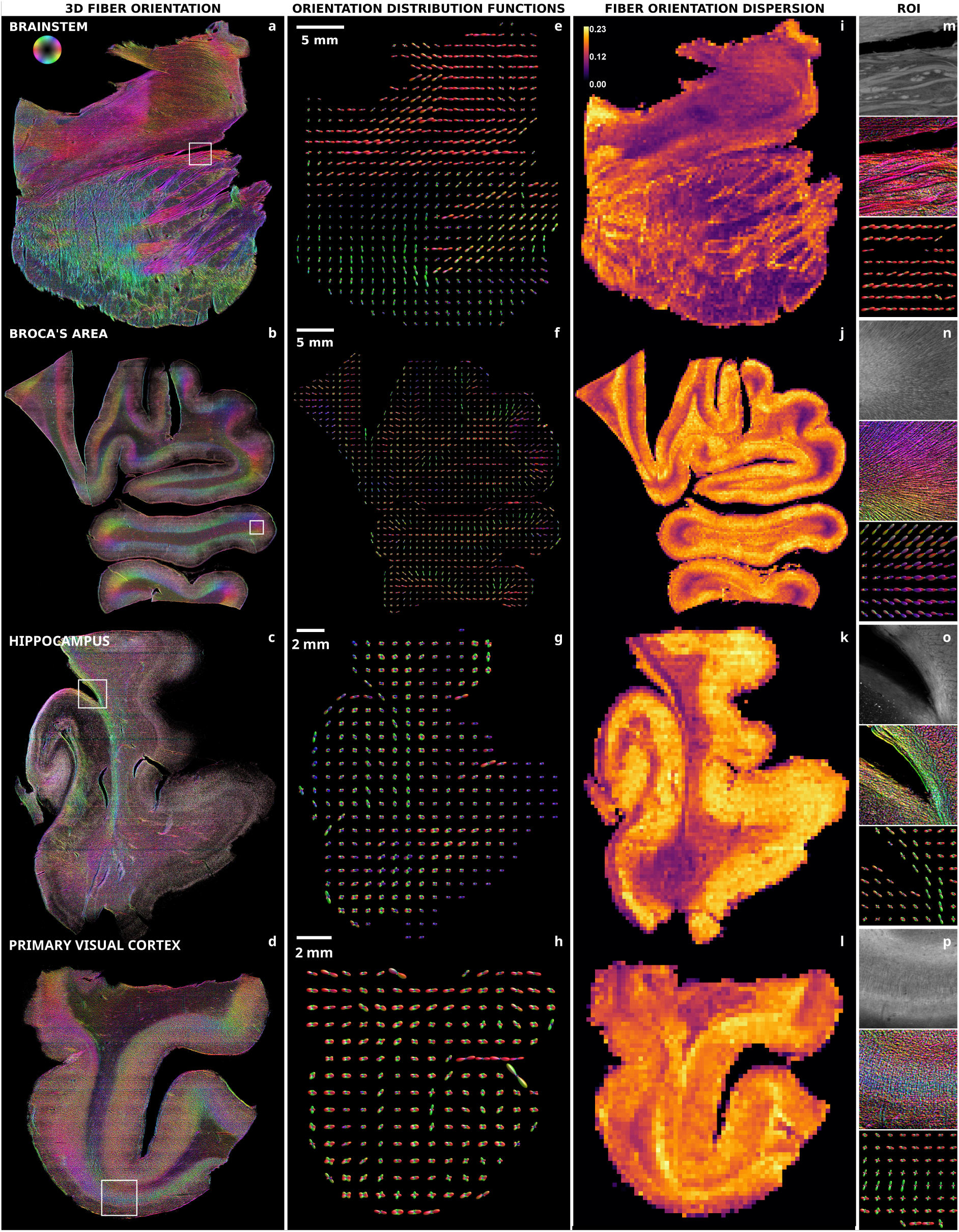
Enhancement and orientation analysis of myelinated fibers in LSFM reconstructions of human brain tissue samples. a-d) Average intensity projections of the 3D fiber orientation HSV colormaps generated by applying Foa3D to the mesoscopic reconstructions of the four anatomical brain regions imaged with LSFM (DiD fluorescence channel, isotropic pixel size: 3.6 ➭m); e-h) ODFs of myelinated fibers (super-voxel size: 1 mm^3^); i-l) orientation dispersion maps of myelinated fibers (isotropic pixel size: 250 ➭m); m-p) 2 mm X 2 mm X 0.25 mm volumetric ROIs (white boxes, ODF super-voxel size: 250 ➭m X 250 ➭m X 250 ➭m).

Notably, fiber bundles extending from the brainstem tissue slab’s locus coeruleus have a precise directionality over their entire length (≈1 cm) (Fig. 3a). Additionally, the dense fiber bundles radiating from the subiculum and the entorhinal cortex, as well as the longitudinal fibers of the alveus and fimbria, have their geometry accurately resolved and characterized (Fig. 3c). Finally, with regard to the spatial orientation colormaps of the Broca’s area and primary visual cortex, Foa3D demonstrates the ability to identify the primarily radial arrangement of myelinated fibers within the gyrus and, on the other hand, the primarily perpendicular orientation to the cortical surface of densely packed fiber bundles (Figs. 3b and 3d).

Furthermore, the ability to faithfully depict the geometry of white matter bundles at larger spatial scales is demonstrated by the maps of orientation distribution functions (ODFs, Fig. 3e-h, super-voxel size: 1 mm^3^) that allow us to highlight the structural coherence across larger regions through the integration of the high-resolution orientation data, providing a valuable link between micrometric details and mesoscopic structural properties. Lastly, the preferential directionality of white matter bundles in relation to the surrounding tissue is successfully revealed by the orientation dispersion maps (OD, Eq. 1, *Fiber orientation analysis (Foa3D)*, *Materials and Methods*; Fig. 3i-l, isotropic pixel size: 250 ➭m). In fact, their near-zero dispersion reinforces the robustness of the Frangi-filter-based analysis by validating the accuracy and consistency of orientation data computed at the voxel level. Together, these fiber orientation images underscore the extreme potential of advanced imaging and image processing tools in unraveling the complexity of the brain’s myeloarchitecture.

## Discussion and Conclusions

In this study, we present a tissue preparation method and an analysis pipeline to investigate the 3D organization of the myelinated fiber architecture in human brain tissues using LSFM (Fig. 4). Our approach was applied to four distinct human brain anatomical regions: the brainstem, hippocampus, Broca’s area and primary visual cortex, demonstrating its versatility. Samples were prepared combining the SHORT clearing and labeling method with a dedicated fiber staining protocol employing the DiD lipophilic dye. Then, we improved the Foa3D image processing tool (see *Fiber orientation analysis (Foa3D)* in *Materials and Methods*) to compute fiber-specific 3D image orientation maps without any human supervision — that is, without creating ground truth fiber orientation data to train a machine-learning-based fiber segmentation algorithm. The ability to perform a fully unsupervised analysis of the myelinated fiber spatial organization, based on a 3D Frangi filter [40], allowed biologists to be relieved from the laborious task of annotating 3D fiber structures in a reference training image dataset. For 3D volumetric reconstruction, we employed a custom-built dual view light-sheet fluorescence microscope [27, 38] enabling micrometer resolution imaging (3.6 ➭m isotropic voxel size) of large tissue volumes (in the cm range) with fast acquisition times (≈0.3 cm^3^/h), providing detailed visualization and distinction of myelinated fiber bundles with challenging geometrical configurations and different cross-sections throughout the entire depth of thick brain tissue slabs.

**Figure 4:**
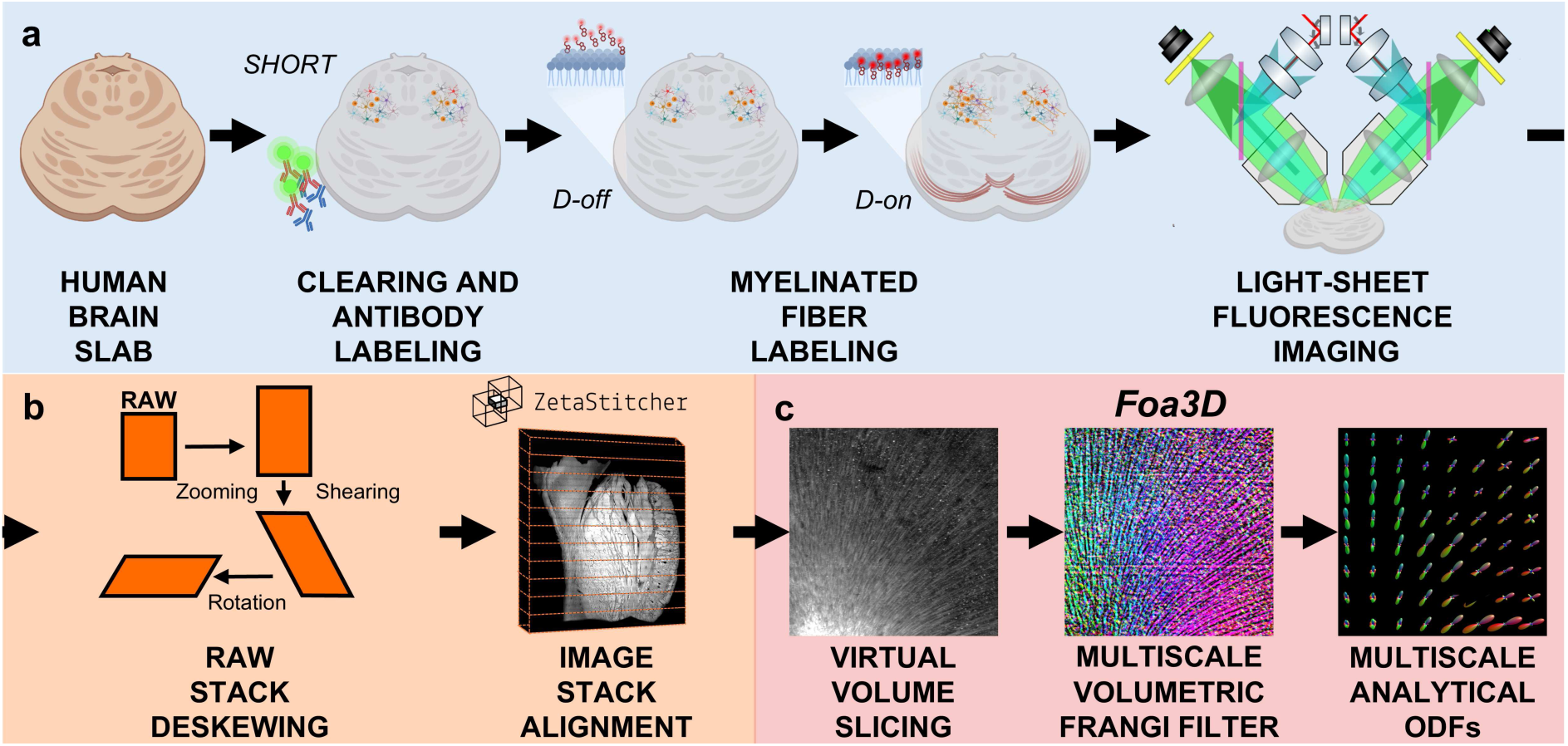
Diagram illustrating the workflow for quantitative 3D myeloarchitectural analysis of human brain tissues. a) Human brain slabs are processed with the SHORT method for tissue delipidation and antibody-mediated labeling of various neuronal populations. Next, in order to stain myelinated fibers, slabs are incubated in D-off solution, to slow the dye’s incorporation into the membrane, followed by incubation in D-on solution to allow DiD molecules to accumulate within the myelin sheath. A custom-built LSFM setup is used for volumetric imaging of the cleared and labeled brain slabs. b) Post-processing of raw LSFM image stacks, which first undergo a composite affine transformation and, then, are automatically aligned using the ZetaStitcher tool for volumetric stitching of large, high-resolution microscopy data. c) Main stages of the unsupervised analysis of 3D fiber orientations performed by the Foa3D Python tool based on Frangi filters.

Compared to other microscopy modalities currently employed in the neuroimaging domain, LSFM possesses unique advantages. In contrast to point scanning techniques such as TPFM, LSFM is characterized by improved volumetric imaging speeds and higher photonic yield, with overall lower light doses to tissue and, thus, decreased adverse photobleaching effects [17]. Compared to the label-free polarimetry-based techniques like PLI and PS-OCT, LSFM achieves higher spatial resolution but requires fluorescent samples. Moreover, a fundamental challenge associated with LSFM imaging is its reliance on tissue transparency. Various methods can achieve optical transparency; however, most involve lipid-removal steps to reduce tissue RI and minimize scattering at water-lipid interfaces, thereby enhancing transparency [14, 15, 16, 18, 41, 42, 43]. This requirement can pose a hurdle when using lipophilic dyes. The multilamellar membrane wrapping that forms the myelin sheath is characterized by a high proportion of lipids (70%–85%) and a low proportion of proteins (15%–30%), in contrast to cell membranes, which have an approximately equal ratio of proteins to lipids [44]. Due to their high affinity for lipid-rich structures, and their cost affordability, lipophilic dyes such as DiI, DiO, Vybrant DiD and FluoroMyelin Green/Red, have been widely used as tracers for neuronal projections in murine tissues for electrophysiological experiments or classical histological preparations [12, 31, 45, 46, 47, 48]. A recent study proposes a near-infrared aggregation-induced emission (AIE)-active probe for myelin imaging also in optically cleared mouse brain slabs [32]. Nevertheless, visualizing the multiscale organization of myelinated fibers deep in human brain tissue requires specific clearing methods that might not always be compatible with dyes that accumulate in lipid-rich structures. In this work, we successfully optimized a myelinated fiber staining approach using the lipophilic DiD dye in combination with the SHORT clearing and antibody labeling method. We also demonstrated that the organic solvent-based methods iDISCO [39], which employ methanol gradients and dichloromethane for lipid removal, could not be successfully combined with a lipophilic-dye staining. We observed a complete absence of fluorescent signals after testing the DiD labeling method on a human brainstem slab cleared with the iDISCO protocol. This may be attributed either to excessive lipid removal or DiD washout following dichloromethane treatment. In contrast, the SHORT tissue transformation method enabled a homogeneous staining of both neuronal bodies, with specific antibodies and myelinated fibers with the DiD probe across multiple thick slabs from the four brain regions investigated, each having a different myelinated fiber content. The SDS-based lipid removal used in this method is milder compared to that of iDISCO, preserving a subset of lipids forming the myelin sheath while still achieving sufficient transparency to reduce light scattering for LSFM imaging. A previous attempt from our group to use the DiD probe in SHORT-processed brain slabs resulted in myelinated fibers being stained to a depth of only 100 ➭m [24]. We improved the penetration rate 5 times by applying DiD through two consecutive 24-hour incubation steps: first, slowing the incorporation of the dye to facilitate its tissue penetration (D-off solution) and then allowing DiD molecules to intercalate into the myelin sheath (D-on solution). This approach prevented the uneven and superficial labeling of myelinated fibers, which is typically caused by the rapid binding of dye molecules relative to their dispersion rate. Notably, the SDS-mediated step required for DiD labeling of fibers did not disrupt prior antibody-based labeling performed with several neuronal markers, including NeuN, somatostatin, and calretinin, nor did it affect the sparse labeling of individual noradrenergic axons with tyrosine hydroxylase. The co-labeling of myelinated fibers and neuronal subpopulations enable correlative studies of myeloand cytoarchitectural features of human brain tissue under both healthy and pathological conditions at the microscale level. Nevertheless, the transparency achieved with SHORT is limited to a thickness of 500 ➭m, while the iDISCO method can clear thicker volumes (up to several mm [49]). Our findings emphasize the importance of selecting and optimizing appropriate tissue-clearing methods for each dye-labeling strategy and application to obtain optimal results.

To analyze the 3D orientation of labeled fibers automatically, we enhanced the Foa3D tool — previously employed on TPFM images [11]— to allow rapid processing of LSFM reconstructions. Such images, in the order of ≈10-1000 GB of data, demand higher computational efficiency to be routinely handled. The implementation of GPU-accelerated fiber enhancement and voxel-level orientation analysis effectively eliminates the critical bottleneck caused by processing the large volumes of data produced by LSFM on a single CPU core. This advancement addresses a major limitation that previously limited this kind of measurement and microscopy modality. Leveraging the unparalleled spatial resolution of fluorescence microscopy, Foa3D can now serve as a powerful tool for obtaining high-resolution fiber orientation information from large tissue samples with minimal computational constraints. The future adoption of advanced rapid image deconvolution and multiview fusion algorithms [50] — suitable for the dual inverted LSFM system utilized in this study — will potentially enable us to enhance the 3D spatial resolution of fiber orientation data to approximately 1 ➭m^3^. Notably, Foa3D’s image processing workflow features the generation of orientation distribution functions (ODFs) at arbitrary spatial scales, which can integrate such fine 3D orientation data at micron resolution to accurately reconstruct the geometry of large fiber bundles. Additionally, the quantification of local orientation dispersion maps enables precise characterization of their preferential directionality relative to the surrounding brain tissue. The improved applicability achieved through our enhancements enables LSFM to be combined with other imaging techniques, such as PLI or PS-OCT, providing complementary 3D fiber orientation data. This integration could be used to validate their orientation estimates or, alternatively, identify potential areas for improvement. Despite the improvements made, Foa3D still has some inherent limitations. Foa3D is designed to analyze the orientation of 3D tubular structures but does not include a stage for the automatic recognition of blood vessels and fibers: this step should be performed separately upstream. Specifically, this could be achieved either by feeding the fluorescence emission channel of myelinated fibers (or vessels) to it, if already highly specific, or, alternatively, by providing the output of a supervised image classifier, such as ilastik [51], if the raw microscopy acquisition contains signals from both structures. Our research group has recently considered, developed, and applied deep-learning-based methods for the automatic detection of neuronal cell bodies and the high-throughput cytoarchitectural analysis of large high-resolution 3D human brain sections (BCFind-v2 [27, 38, 52]). For this task, more than 20,000 cell centroids had to be manually tagged in order to train a U-Net-based 3D cell body detector with adequate performance on unseen brain tissue reconstructions. However, we deemed this considerable manual annotation effort to be undoubtedly lower with respect to the hypothetical creation of fiber orientation ground truth data by sparse manual tracing, that a robust model for single-fiber or fiber bundle segmentation would require to be trained, making this approach currently unfeasible. Nevertheless, in the future, researchers may look into using micrometer-resolution 3D fiber orientation data automatically generated by Foa3D, and properly validated by human inspection, to help train a generative deep learning model for fiber tracing and orientation analysis in large brain tissue datasets obtained by LSFM. In conclusion, establishing a comprehensive map of the mammalian brain connectome remains limited by methodological barriers related to assess and measure connectivity and its complexity across scales, even in small model species such as the mouse. As such, available methods for mapping brain connectivity yield incomplete datasets leading to inaccuracy when applying them to larger species, particularly nonhuman primates and humans, or to in vivo investigations. Such methods have a clear potential in the study of brain health and brain disorders. Vulnerability of select brain areas, neuronal subpopulations, and connections is known to occur in many neuropsychiatric diseases. The pathogenic mechanisms involving specific cell types and states, cell-cell interactions, and neural pathways, as well as the role of signaling and spatial interactions with non-neuronal cell types, are emerging from single-cell and spatial approaches in many age-related neurodegenerative disorders and neurodevelopmental illnesses [53, 54, 55, 56, 57]. In this context, the innovative fiber staining method based on DiD, combined with the multiscale fiber dispersion maps generated by the Foa3D tool, may provide a quantitative framework for assessing the human brain connectome. Its broad applicability could be also used to investigate pathological alterations, offering valuable insights into connectivity disruptions and abnormal remodeling of the white matter architecture in disease states.

## Materials and Methods

### Samples

The human brain samples used in this paper were obtained from control subjects who died of natural causes with no clinical diagnoses or neuropathology. A standard fixation protocol was used in which the brain was fixed in 10% formalin for a minimum of 90 days. Hippocampus and primary visual cortex brain samples were obtained from the body donation program (Association des dons du corps) of Université de Tours, France. Prior to death, participants gave their written consent for using their entire body — including the brain — for any educational or research purpose in which anatomy laboratory is involved. The authorization documents (under the form of handwritten testaments) are kept in the files of the Body Donation Program. Brainstem and Broca’s area samples were procured from the Department of Neuropathology at the Massachusetts General Hospital Autopsy Service (Boston, USA). Written consent was obtained from healthy participants before death, following institutional review board–approved tissue collection protocols from Partners Institutional Biosafety Committee (protocol 2003P001937). Tissue sections were obtained using a custom-made vibratome (Compresstome VF-900-0Z) and stored at 4 ^◦^C in PBS 0.02% NaN_3_.

### Tissue preparation

#### SHORT tissue transformation method

Slabs with a thickness of 300 ➭m or 500 ➭m were processed using the SHORT method, as previously described [24, 25]. The specimens were first incubated in a Switch-Off solution (50% v/v 1X PBS (pH 3), 25% v/v 0.1 M hydrochloric acid (HCl), 25% v/v 0.1 M potassium hydrogen phthalate (KHP), and fresh 4% v/v glutaraldehyde) and, after 24 h, this was replaced with the Switch-On solution (1X PBS (pH 7.4) and fresh 1% v/v glutaraldehyde). Both incubations were performed at 4 ^◦^C with gentle shaking. Afterwards, the slabs were washed three times in 1X PBS at room temperature (RT), followed by incubation in inactivation solution (1X PBS (pH 7.4), 4% w/v acetamide, 4% w/v glycine, pH 9.0) at 37 ^◦^C. Following the inactivation of the reactive glutaraldehyde, the samples were washed three times in 1X PBS at RT, with each wash lasting 2 h, and then incubated in the clearing solution (200 mM sodium dodecyl sulfate (SDS), 20 mM sodium sulfite (Na_2_SO_3_), and 20 mM boric acid (H_3_BO_3_), pH 9.0) in a water bath at 55 ^◦^C for 6 days. After the clearing step, the samples were extensively washed in 1X PBS with 0.1% Triton X-100 (PBST) for 24 h with gentle shaking. Quenching of the autofluorescence was performed by treating the slabs with 30% H_2_O_2_ for 90 min at RT, followed by antigen unmasking through incubation in a preheated antigen retrieval solution (10 mM Tris base (v/v), 1 mM EDTA solution (w/v), 0.05% Tween 20 (v/v), pH 9) at 95 ^◦^C in a water bath for 10 min. After cooling to RT, the specimens were washed in deionized water for 5 min and then equilibrated in PBS 1X for 1 h. The specimens were then incubated with primary antibodies diluted in PBST with 0.01% w/v NaN_3_, at 37 ^◦^C with gentle shaking for 5 days. The primary antibodies used are: anti-NeuN antibody (Merck Life Science, #ABN91; RRID:AB 11205760), diluted 1:50; anti-tyrosine hydroxylase (Merck Life Science, #AB152; RRID:AB 390204), diluted 1:500; anti-somatostatin (Santa Cruz Biotechnology, #sc.47706; RRID:AB 628268) diluted 1:200; and anti-calretinin (Protein-Tech, 12278-1-A; RRID:AB 2228338) diluted 1:200. After 24 h of washing with PBST at 37 ^◦^C, the stained samples were incubated with the secondary antibody diluted 1:200 in PBST with 0.01% w/v NaN_3_ at 37 ^◦^C with gentle shaking for 5 days. The secondary antibodies used are: Goat anti-Chicken IgY (H+L) Alexa Fluor 647 (Invitrogen, #ab150171, RRID:AB 2921318); Goat anti-Rabbit IgG (H&L) Alexa Fluor 568 (Abcam, #ab175471, RRID:AB 2576207); Donkey Anti-Rat IgG (H&L) Alexa Fluor 568 (Abcam, #ab175475, RRID:AB 2636887); Alpaca Anti-Rabbit IgG (H&L) Alexa Fluor 488 (Jackson ImmunoResearch Labs; #611-545-215, RRID:AB 2721874).

#### iDISCO method

A 300 ➭m-thick slice was processed using a modified iDISCO protocol [39]. The specimens were first dehydrated in increasing concentrations of methanol (MeOH)/H_2_O (20, 40, 60, 80, and 100%) with gentle rotation for 30 min each at RT. The slabs were then incubated overnight at RT with gentle rotation in a 66% dichloromethane (DCM)/33% MeOH solution. Afterwards, the samples were washed twice with 100% MeOH for 30 min. Autofluorescence was then reduced by incubating the samples in 5% H_2_O_2_ in MeOH at 4 ^◦^C overnight without shaking. The following day, the samples were rehydrated using decreasing concentrations of MeOH/H_2_O (80, 60, 40, and 20%), followed by one wash in PBS 1X and two in PBST, all performed with gentle rotation for 30 min each at RT. On the same day, the samples were incubated in a permeabilization solution (1X PBS, 0.2% v/v Triton X-100, 20% v/v DMSO and 0.3 M glycine), overnight at 37 ^◦^C with gentle rotation. The next day, they were incubated in a blocking solution (1X PBS, 0.2% v/v Triton X-100, 0.2% w/v gelatin porcin skin and 0.01% w/v NaN3), for 6 h at 37 ^◦^C with gentle rotation. The fiber staining with the DiD dye was performed at this point, following the protocol described in the section below. After staining, slabs were washed once in PBSTw-Heparin (1X PBS, 0.2% v/v TWEEN-20 and 0.01 mg/mL Heparin), dehydrated as previously, described and incubated for 3 h in 66% DCM/ 33% MeOH. Finally, the specimens were incubated twice with 100% DCM for 20 min, followed by RI matching in dibenzyl ether (DBE) at RT.

#### DiD staining protocol

After clearing and antibody labeling using either SHORT or iDISCO protocol, myelinated fibers were stained using the lipophilic carbocyanine dye DiD solid (Invitrogen, #D7757). A stock solution (3 mg/mL) was prepared by dissolving lyophilized DiD in a solution containing 10% SDS in 1X PBS, which was then stored at RT. Tissue slabs were first equilibrated in a solution of 10 mM SDS in 1X PBS at 37 ^◦^C with gentle shaking overnight. The following morning, they were incubated in the D-off solution (0.025 mg/mL DiD, 10 mM SDS in 1X PBS) for 24 h at 37 ^◦^C with gentle shaking, followed by an additional 24-hour incubation in the D-on solution (0.025 mg/mL DiD, 1X PBS with 0.1% Triton X100) under the same conditions. To remove the unbound dye, the slabs were extensively washed in 1X PBST at 37 ^◦^C with gentle shaking for 24 h and RI matching was performed based on the protocol used.

### Imaging

#### Spinning disk microscope

Images of human brainstem slabs in Fig. 1 were acquired using a Nikon confocal spinning disk microscopy (Nikon Ti2-CrestV3) and processed using the NIS-Elements AR 5.42.06. The following objectives were used: Nikon CFI Plan Apo 10XC GLYC, NA: 0.5, WD: 5.5 (material number: MRD71120); CFI Plan Fluor 20XC MI, NA: 0.75, WD: 0.51-0.33 (material number: MRH07241); CFI Plan Apochromat Lambda D 60X Oil, NA: 1.42, WD: 0.15 (material number: MRD71670). The laser sources have nominal peak wavelengths of 488 nm, 561 nm and 640 nm, while fluorescence emission was detected using single-bandpass filters at 525/45 nm, 609/54 nm and 698/70 nm, respectively.

#### Light-sheet fluorescence microscope

The brain tissue samples were imaged using a specially designed inverted LSFM system that was detailed in one of our group’s earlier studies [26, 27, 38]. This custom LSFM setup includes two identical 12X objectives (LaVision Biotec, Germany, LVMI-Fluor 12X PLAN NA: 0.53, WD: 8.5 to 11 mm). These are oriented at 90^◦^ relative to each other and spaced so that their fields of view (FOVs) overlap in the center. The objectives are inclined at 45^◦^ with respect to the horizontal plane, where the sample holder is positioned, and they alternately play excitation and detection roles. The microscope is equipped with four laser diode modules (Cobolt, Sweden) having the following wavelengths and nominal maximal output powers: (405 nm, 100 mW); (488 nm, 60 mW); (561 nm, 100 mW); and (638 nm, 180 mW). The four gaussian beams are combined by means of three dichroic mirrors after being magnified by dedicated telescopes. Next, the resulting combined beam is divided by a 50 to 50% beam splitter and outgoing light is then addressed in two identical excitation pathways, each comprising an acousto-optical tunable filter (AOTF, AOTFnC-400.650-TN, AA Optoelectronic, France) for beam intensity, phase, and wavelength modulation. The two AOTFs also enable the independent shuttering of each illumination pathway thus preventing the introduction of stray light from the excitation beam during image acquisition. The planar light-sheet illumination is created by beam-scanning through a galvo mirror (6220H, Cambridge Technology, USA) and subsequent focusing into a scan lens (#45-353, Edmund Optics, USA; focal length (FL) = 100 mm, achromat), which converts the angular deflection of the galvo scanner into a lateral displacement of the incident light, resulting in a digitally-scanned selective illumination of a single layer of the tissue. The scan lens is placed in a plane conjugated with the back focal plane of the objectives, which are in turn mounted on motorized stages (PI L-509.14AD00, Physik Instrumente, Germany) to allow for focus adjustment. A 3-axis motorized stage system (two PI M-531.DDG and a PI L-310.2ASD, Physik Instrumente; overall motion range: 30 cm X 30 cm X 2.5 cm) is used to translate the sample, that is imaged while being shifted along the horizontal x-axis through the fixed FOVs of the two objectives. A multiband dichroic beamsplitter (Di03-R405/488/561/635-t3-55x75, Semrock, USA) is used to separate the reflected excitation light from the fluorescence emitted by the sample before the detection tube lens (#45-179, Edmund Optics, US; FL = 200 mm, achromat) directs it onto a sCMOS camera (Orca-Flash4.0 v3 Hamamatsu, Japan; pixel size: 6.5 ➭m X 6.5 ➭m, pixel array: 2048 X 2048, 16 bit). To selectively image the variously labeled structures within the brain tissue samples, four different bandpass filters were placed in front of the cameras on motorized filter wheels (FW102C, Thorlabs, USA). By synchronizing the rolling shutter sweep with the galvo scan of the light sheet, cameras operate in confocal detection mode [58]. The excitation and detection modes of each optical route are separated by a half-frame time delay to prevent excitation stray light from interfering with the active rolling-shutter rows on the recording camera. The tissue sample is moved along the y-axis (by a range below the FOV side: ≈1109 ➭m) once an image stack is finished, and the process is repeated until the entire tissue volume has been acquired. For an image overlap larger than 100 ➭m, a y-step of 1 mm was used.

### Image processing

#### LSFM image post-processing

In order to obtain isotropic volumetric images correctly aligned with the sample holder, LSFM image stacks had to be properly processed because both the cameras and the objectives of the custom-made LSFM setup are inclined by 45^◦^ with respect to the sample plane. Therefore, by means of a custom Python tool, raw stacks underwent a composite affine transformation (scaling, shearing, and rotation) generating images oriented according to the laboratory’s reference system, with a final isotropic voxel size of 3.6 ➭m. Then, using the open-source ZetaStitcher tool (https://github.com/lens-biophotonics/ZetaStitcher), adjacent, overlapping stacks of LSFM images were aligned and finally fused [59]. By analyzing the FFT-based cross-correlation of overlapping regions at specific depths, ZetaStitcher effectively determines the best alignment of neighboring picture tiles, allowing for the high-throughput stitching of sizable volumetric microscopy datasets. For neighboring image stacks obtained with the employed LSFM setup, the nominal vertical tile overlap of ≈100 ➭m ensured that no spurious doubling of related tissue structures occurred inside matching tissue areas.

Fiber orientation analysis (Foa3D)

An upgraded version of Foa3D (3D Fiber Orientation Analysis, https://github.com/lens-biophotonics/Foa3D), first presented in Sorelli M. et al., 2023 [11] was used to compute 3D fiber orientations within the four anatomical regions of the human brain imaged with LSFM. Foa3D is a Python tool leveraging multiscale 3D Frangi filters [40] for obtaining a ground-truth-free enhancement of tubular structures of varying diameters and, thus, a specific evaluation of the orientations of single neuronal fibers (e.g., within grey matter) and/or tightly packed white fiber tracts. Exploiting ZetaStitcher’s API for querying arbitrary subvolumes of tiled tissue reconstructions, Foa3D supports mesoscopic micrometer-resolution LSFM images that exceed the memory typically available on low-resource machines.

The Frangi filter’s response is affected by a range of scale and sensitivity parameters that need careful tuning in order to achieve the desired enhancement of elongated fiber structures. As previously discussed [11], the optimal scale that best preserves the intensity and cross-sectional size of a 3D tubular structure corresponds to half of its expected radius. In the present work, two separate scale values, namely 5 and 10 ➭m, were adopted for detecting fibers having nominal diameters of 20 and 40 ➭m (i.e., more than 5 pixel wide). Furthermore, the three sensitivity parameters of the Frangi filter (α, β, γ) were systematically optimized to enhance the selective detection of fiber structures in LSFM image reconstructions. Initially, the α and β sensitivity parameters, associated with local geometrical grey-level-invariant features (i.e., image plateness and blobness, respectively), were adjusted through a systematic 2D grid search, sampling four values over a logarithmic scale (α, β ∈ [10^−3^, 10^0^]). The response of the Frangi filter was qualitatively evaluated within a tissue region of interest (ROI) extracted from the Broca’s area reconstruction, including distinct and clearly recognizable fiber bundles. During this stage, the background contrast sensitivity (γ) was kept constant at 1000. As a result of this process, the sensitivity configuration α = 0.001 and β = 1 was identified as the most effective, providing the highest selective enhancement of fiber structures while effectively suppressing contributions from the neuronal cell soma. Subsequently, the γ sensitivity parameter was refined by testing four values over a logarithmic scale (γ ∈ [100, 103]) to further optimize the filter’s overall response based on the specific greyscale range of each LSFM reconstruction. This adjustment revealed that γ = 100 was the most effective value for the tested ROI, achieving a balance between enhancing fiber structures and minimizing background noise. To prevent undesired local discontinuities between the fiber orientation fields derived from adjacent tiles, a global, spatially invariant contrast sensitivity was adopted instead of automatically adjusting γ based on the local Hessian norm within each subvolume, as originally suggested by Frangi et al. [40]. To mask dependable fiber structures and generate fiber-specific orientations, a global threshold of 0.9 was consistently applied to the Frangi filter’s response (comprised between 0 and 1) across the LSFM image dataset. Again, automatic thresholding algorithms working on separate subvolumes were avoided at this stage for preventing local tiling artifacts. High-resolution fiber orientation data computed at the native voxel size of the LSFM reconstructions were then integrated using the fast analytical approach proposed by Alimi et al. [60] into orientation distribution functions (ODFs), providing a comprehensive characterization of 3D fiber tract orientations within larger spatial compartments, as previously shown [11]. Within such compartments, furthermore, a normalized index of fiber orientation dispersion (OD) was quantified from the eigenvalues of the local orientation tensor — defined as the sum of the outer products of the 3D fiber orientation vectors — which captures the dominant local orientations and their distributions, as follows:

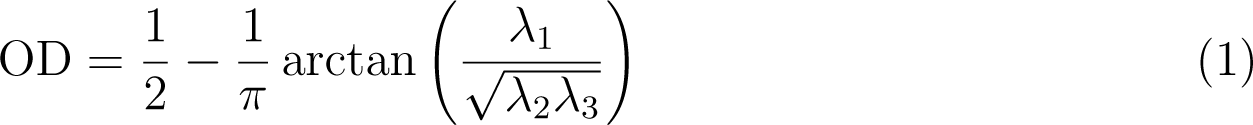

where λ*_i_* are the eigenvalues of the fiber orientation tensor in ascending order. The entire image processing pipeline was executed on a high-throughput computing cluster consisting of four nodes managed using the HTCondor Software Suite [61]. The computational performance of the original Foa3D tool, which runs on a single CPU, was compared to that of the new version incorporating a GPU-accelerated Frangi filtering stage. Specifically, the original version was evaluated on two identical nodes, each equipped with two Intel^➤^ Xeon^➤^ Silver 4214R 12-core processors (2.4 GHz) and 192 GB of RAM. Instead, the new version was assessed on a separate node featuring an NVIDIA GeForce RTX➋ 3090 GPU with 24 GB of G6X memory.

## Code Availability

The new Foa3D featuring GPU acceleration will be released upon paper acceptance at https://github.com/lens-biophotonics/Foa3D. The ZetaStitcher software is freely available at https://github.com/lens-biophotonics/ZetaStitcher. The SPIMlab tool is available at https://github.com/lens-biophotonics/SPIMlab.

## Data Availability

The datasets generated and analyzed during the current study are available from the corresponding author on reasonable request.

## Acknowledgements

We express our gratitude to the donor involved in the body donation program of the Association des dons du corps du Centre Ouest, Tours, France, and Massachusetts General Hospital Autopsy Service, USA, who made this study possible by generously donating their body to science.

## Author Contributions

MS developed Foa3D and conducted the 3D fiber orientation analysis of the microscopy tissue reconstructions. DDM and IC optimized the clearing method and the labeling of myelinated fibers. SB contributed to and tested the GPU-accelerated version of Foa3D. DDM and JR prepared the tissue samples. DDM, LP, and FC performed light-sheet microscopy imaging. MS, SB, and GM carried out microscopy image postprocessing for the generation of 3D mesoscopic tissue reconstructions. FSP and IC acquired financial support and FSP provided the equipment employed in this work. PRH provided neuroanatomical expertise. CD dissected and fixed human brain samples. IC conceived and supervised the study. MS, DDM, and IC interpreted the results and wrote the manuscript with input from all the authors.

## Supplementary Material

**Figure S1:**
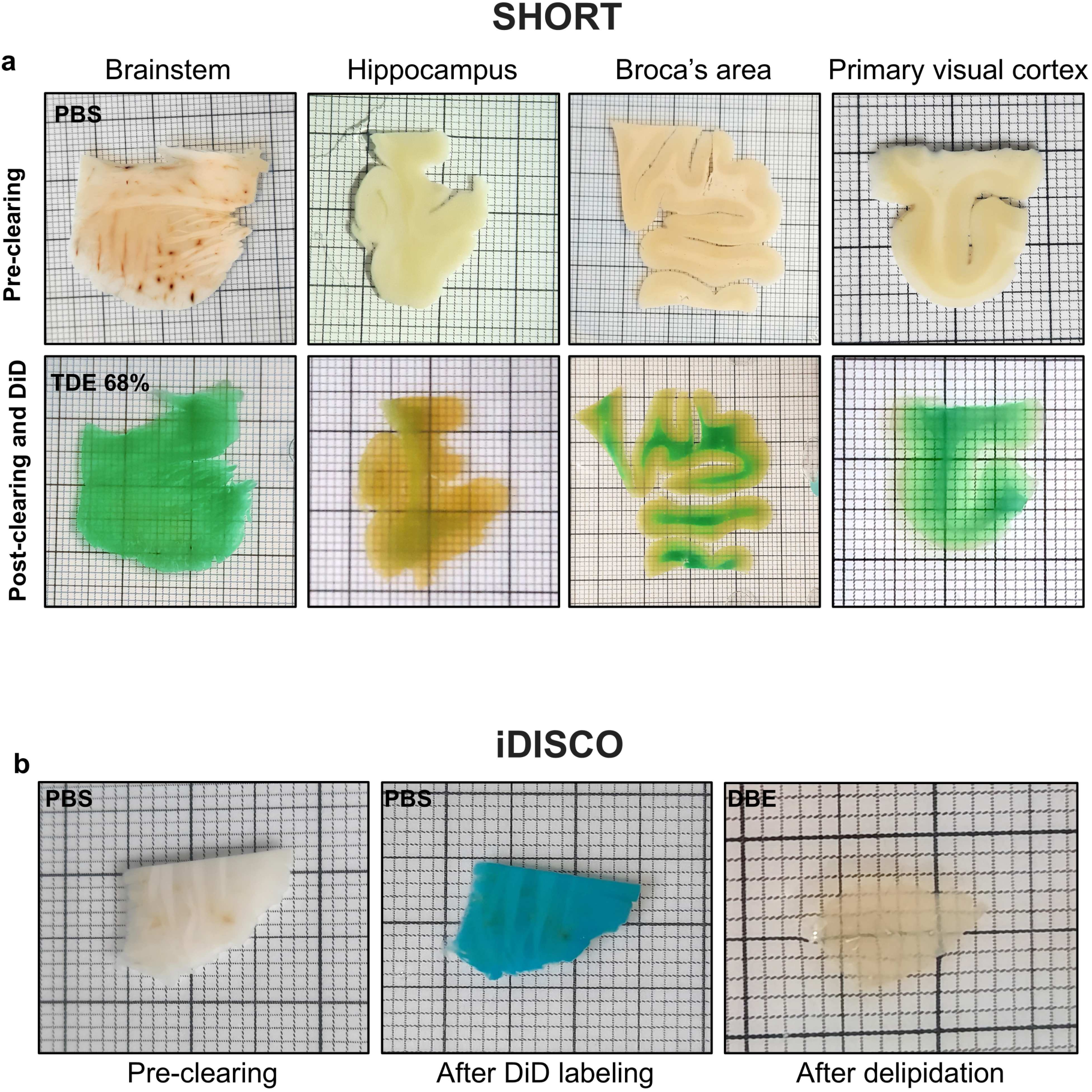
Pre- and post-clearing images of multiple human brain slabs. a) Images showing the different slabs before the whole process (top: PBS) and after clearing, labeling and refractive index (RI) matching using the SHORT method combined with the DiD protocol for myelinated fiber labeling (bottom: TDE 68%). b) Images showing a brainstem slab portion before clearing (left), after the application of the DiD dye (center), and after delipidation using the iDISCO protocol and RI matching in DBE (right).

**Figure S2:**
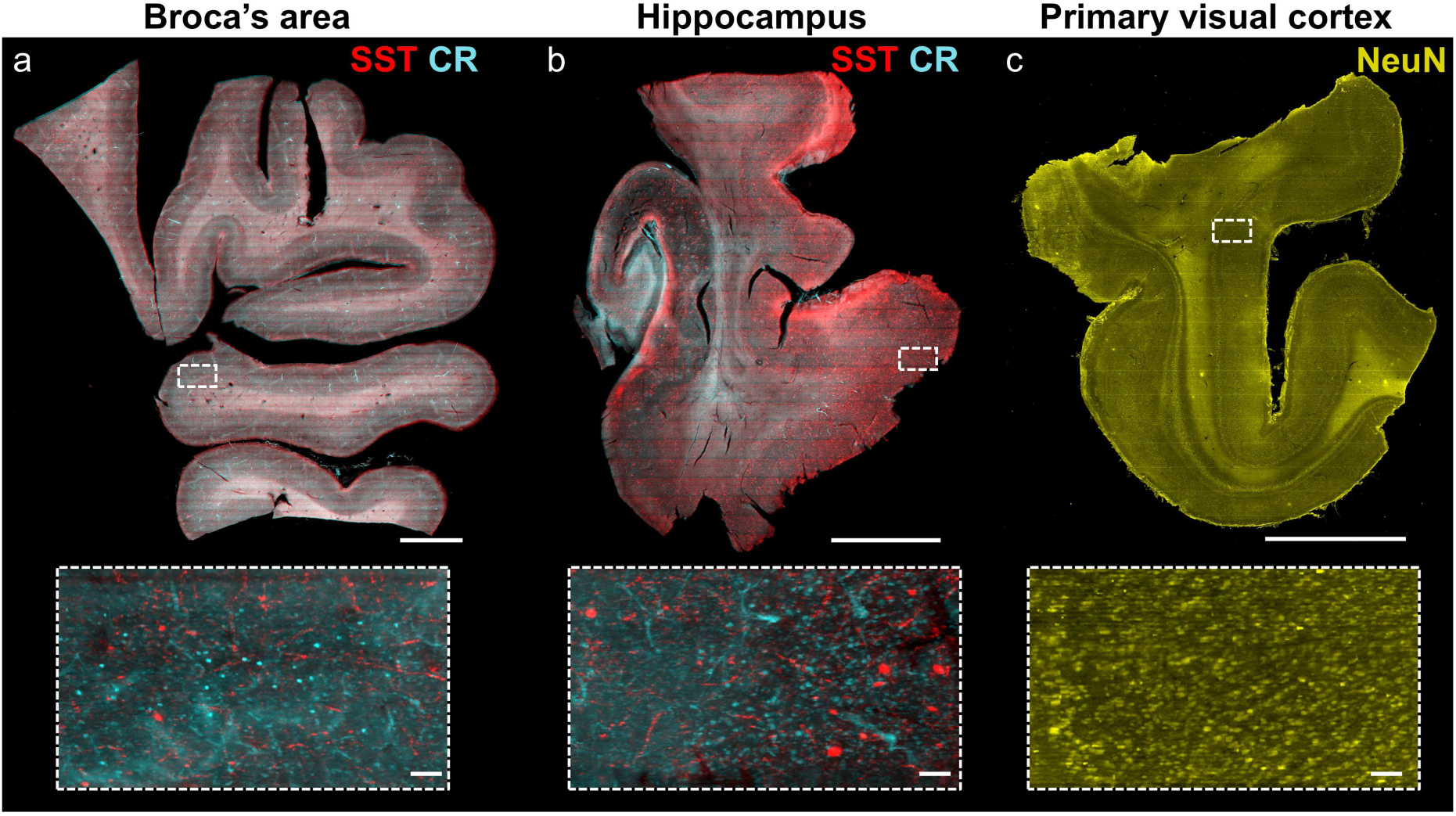
Antibody co-labeling and LSFM imaging of multiple subpopulations of neurons in human brain slabs. Multichannel maximum intensity projection images (3.6 ➭m isotropic voxel size) of SHORT-processed slabs. a) 500 ➭m-thick adult postmortem human Broca’s area labeled with anti-somatostatin (SST, red) and anti-calretinin (CR, cyan) antibodies. Scale bar: 0.5 cm. b) 500 ➭m-thick adult postmortem human hippocampus labeled with anti-somatostatin (SST, red) and anti-calretinin (CR, cyan) antibodies. Scale bar: 0.5 cm. c) 500 ➭m-thick postmortem human primary visual cortex labeled with anti-NeuN antibody. Scale bar: 0.5 cm. Magnified insets are shown at the bottom and identified by white boxes in the upper images. Scale bar: 100 ➭m.

